# Abrupt transitions and its indicators in mutualistic meta-networks: effects of network topology, size of metacommunities and species dispersal

**DOI:** 10.1101/2022.05.02.490298

**Authors:** Gaurav Baruah

## Abstract

Gradual changes in the environment could cause dynamical ecological networks to abruptly shift from one state to an alternative state. When this happens ecosystem functions and services provided by ecological networks get disrupted. We, however, know very little about how the topology of such interaction networks can play a role in the transition of ecological networks at spatial scales. In the event of such unwanted transitions, little is known about how statistical metrics used to inform such impending transitions, measured at the species-level or at the community-level could relate to network architecture and the scale of spatial interactions such as the size of the metacommunity. Here, using hundred and one empirical plant-pollinator networks in a spatial setting, I evaluated the impact of network topology and spatial scale of species interactions on abrupt transitions, and on statistical metrics used as predictors to forecast such abrupt transitions. Using generalized Lotka-Volterra equations in a meta-network framework, I show that species dispersal rate and the size of the metacommunity can impact when an abrupt transition can occur. In addition, forecasting such unwanted abrupt transitions of meta-networks using statistical metrics of instability was also consequently dependent on the topology of the network, species dispersal rate, and the size of the metacommunity. The results indicated that the plant-pollinator meta-networks that could exhibit stronger statistical signals before collapse than others were dependent on their network architecture and on the spatial scale of species interactions.

## Introduction

Ecological systems could abruptly shift from one state to another in response to gradual changes in environmental conditions. Such abrupt transition occurs when environmental conditions cross a specific threshold, the threshold being commonly known as a tipping point (Scheffer et al. 2001; Dakos et al. 2014). Tipping points and abrupt transitions are generally observed in ecological systems governed by positive feedback loops. Abrupt transition can occur in systems ranging from acquatic systems such as algae and macrophyptes (Dakos et al. 2018), mutualistic ecological networks (Dakos and Bascompte 2014; Lever et al. 2014; Baruah 2022), and populations exhibiting allee thresholds (Hilker 2010). When such abrupt transition occur, the equilibrium state of the system jumps to another state where loss of beneficial ecosystem functions and services could occur (Hutchings and Reynolds 2004; Scheffer 2009; Dunne and Williams 2009). Abrupt transitions, thus, can cause long-term radical changes to ecosystems. However, at the scale of multiple communities connected by species dispersal, very few studies have explored how local network architecture and spatial scale of interactions, such as size of the metacommunity, could impact the timing of abrupt transitions .

Mutualistic communities are examples of communities that exhibit positive feedback loops in networks of interactions between two groups of species (Bascompte and Jordano 2013; Dakos and Bascompte 2014; Kéfi et al. 2016; Metelmann et al. 2020). While negative interactions such as intraspecific competition could have stabilising effects, positive interactions such as those observed in plant-pollinator or mutualistic networks, could be destabilising and could result in the presence of alternative stable states (Kéfi et al. 2016; Baruah 2022). The spatial context of such transition has somewhat remain unexplored, particularly on the impact of topological and architectural aspects of such systems such as network size, nestedness, or connectance. Nestedness is an ecological pattern that has been widely reported for species occurrences as well as in species interaction networks (Bascompte and Jordano 2013). Nestedness occurs when specialist species in an ecological network interact more with subsets of species that interact with generalist species . Connectance, on the other hand, describes the realised number of species interactions in a community out of all possible interactions. These two topological features capture different aspects of species interaction networks and has been suggested to impact stability of communities. However, their impact on tipping points in a spatial context remain somewhat unexplored (but see Revilla et al. (2015)). Previous studies on spatial mutualistic systems has suggested that the presence of transitions can be modulated by shift in strength in species phenology or habitat destruction (Fortuna and Bascompte 2006). Habitat destruction increases the chances of abrupt transition which rewires local networks (Revilla et al. 2015). In addition, with strong mutualistic interactions the amount of habitats required for persistence of all species then depends on the range of species dispersal (Prakash and Roos 2004). However, it is somewhat unknown whether topological features of mutualistic networks such as network size or connectance could influence the timing of occurrence of abrupt transition in response to gradual changes in environment (Metelmann et al. 2020), especially when different spatial scale of interactions could be at play.

Rarely, ecologlical communities occur in isolation. More often, ecological communities occur in habitat patches across a larger network of patches that is connected by dispersal, which is commonly known as a metacommunity. Metacommunity concept has gained much attention over the last decade and empirical and theoretical studies have provided with an understanding of how local and regional processes work to maintain diversity both at local and spatial scale (Loreau 1998; Limberger et al. 2019). Species diversity can stabilize the dynamics of local communities, and species dispersal can provide spatial insurance. This stabilization in local dynamics can occur due to different asynchronous responses of species over time to temporal changes in the environment (Loreau et al. 2003; Heino et al. 2015; Shoemaker and Melbourne 2016). On the other hand, spatial insurance in metacommunities arises when local communities exhibit asynchronous dynamics. This happens when species dispersal rate is limited or when species composition across local communities vary considerably thereby leading to spatial heterogeneity (Wang and Loreau 2014; Walter et al. 2017). Species dispersal from different habitat patches could potentially rescue local communities from collapses as environment changes. However, the role of species dispersal on timing of transition of meta-networks remains unknown. For instance, does spatial scale of species interactions and the spread of such ecological networks (number of habitat patches) determine whether network collapses occur earlier or later?

Local and regional scale extinctions could occur not only due to local processes such as predation or competition but also due to large scale external disturbances (Cunillera-Montcusí et al. 2021). Large scale disturbances akin to changes in climate could impact species interactions not only locally but also across communities connected in space (Revilla et al. 2015; Morton and Rafferty 2017; Thompson and Gonzalez 2017; Renner and Zohner 2018; Kudo and Cooper 2019). However, such species mismatches could be rescued when similar communities are accessible to species in a mosaic of communities connected by species dispersal. It is, however, unknown whether such global changes in the environment interacts with local network topological properties to mitigate drastic change. Increases in phenological mismatch could be further compounded by habitat destruction and could lead to substantial changes in network architecture (Revilla et al. 2015). In addition, whether the spatial scale (size of the metacommunity) of such changes matters in the occurrence of transition is somewhat not known.

There has been statistical tools developed to inform impending transitions that could occur as environment gradually changes (Scheffer 2009). Abrupt transitions could occur when changes in the environment crosses a certain threshold that pushes the ecological system towards another alternative state where ecosystem functions could be lost permanently. However, there are statistical tools that been developed to forecast such impending transitions which are commonly known as “early warning signals”. Commonly used signals are temporal autocorrelation and variance that could derived using a sliding window approach (see Dakos et al. (2012a) for details) from state based temporal data such as abundance or biomass. However, the utility of such signals are dependent on a host of factors that includes sampling requirements (Arkilanian et al. 2020), data quality (Clements et al. 2015b), eco-evolutionary factors (Baruah et al. 2020, 2021) and type of species interactions (Dakos 2017; Patterson et al. 2021; Baruah et al. 2022). One important challenge is to test the utility of such signals in a multispecies context embedded within a spatial scale of species interactions. This is especially relevant as multispecies communities rarely occur in isolation and are generally connected by dispersal among habitat patches. The detection of such signals becomes even more challenging as dynamics of such communities are inherently linked to their topological features which also directly impacts the occurrence of transitions (Dakos and Bascompte 2014; Baruah 2022).

Here, using hundred and one empirical plant-pollinator networks in a spatial context collated from web-of-life.es database, I explore how the effects of topological network features such as network size, connectance, or nestedness can interact with spatial scale i.e., size of the metacommunity ( number of habitat patches i.e., 2, 5 ,10, 20) to impact the timing of transitions and on indicators of temporal and spatial resilience. Using generalized Lotka-Voltterra equations, I model the ecological dynamics of spatially-explicit mutualistic meta-networks to global changes in strength in mutualistic interactions. Specifically, using different metacommunity sizes, I show that timing of abrupt transition depends not only on how species disperse across habitat patches, but also on local network topological predictors such as network size and connectance of the network. Furthermore, the threshold at which a species could abruptly transition is influenced by the degree of the species, rate of dispersal and also by the size of the metacommunity. In addition, how large the metacommunity was, also significantly played a role on how early an abrupt transition might occur. Furthermore, when such global transition of mutualistic metacommunities occur, predictability with temporal and spatial resilience indicators also depends on the topological network features and on the rate of dispersal of species. These results argue the importance of understanding the dynamics of communities from a spatial perspective and highlights the importance of network architecture on biodiversity maintenance.

## Methods

Using www.web-of-life.es database, I collated hundred and one empirical plant-pollinator networks (see table S1 in supplementary appendix 2 for details and the references). I chose hundred and one networks to ensure that I had sampled networks that display a wide range of topological properties. These empirical networks were set up in a spatially-explicit landscape of different sizes of two, five, ten, and twenty habitat patches that determined the spatial scale of mutualistic interactions. This spatial scale determined to what extent plant-pollinator interactions were impacted when global changes occur at the scale of the metacommunity. For instance, when changes in climate that drives changes in phenological interactions occur, does spatial scale of mutualistic interactions matter in delaying an abrupt transition to collapse? To be noted that this study is purely exploratory and theoretical in nature. The only empirical part of this study comes from the fact that the networks used in simulating the dynamics were collected from field studies.

These spatially explicit landscape were set up in a two dimensional landscape (see Grilli et al. 2015 for details). All habitat patches were connected. Following this, I model the ecological dynamics of mutualistic interactions using generalized Lotka-Volterra equations:

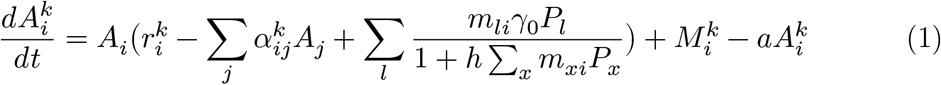

Similarly, the dynamics of plants on a spatially explicit metacommunity can be written as:

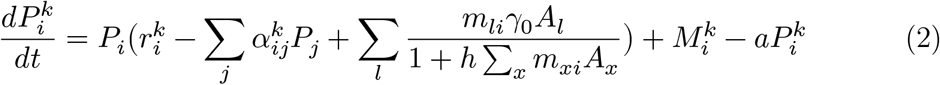

where 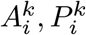 represents pollinator and plant abundance for species *i* in habitat patch 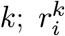 is the species specific growth rate independent of mutualistic interactions at patch 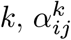 represents interspecific competition within each guild of species at patch *k*, *γ*_0_ represents the average mutualistic interaction strength when a plant and a pollinator interacts, with *m_li_* determining the network structure and is either 0 or 1 depending on whether an interaction exist between a plant and a pollinator, *h* is the handling time which was fixed at 0.15, which denotes the amount of time a pollinator takes to interact with a plant. Handling time was fixed at 0.15 as it falls within the range observed in empirical plant-pollinator interactions (see Klumpers et al. (2019)). 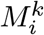 is the density of species that arrives from all the patches in the metacommunity, and finally a fraction of individuals emigrates from a habitat patch *k* at the rate *a*.

In the model simulations, I ensured that mutualism was obligate without any loss of generality. This meant that growth rates of species 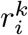 was negative for both the guilds of species and species persistence was dependent on mutualistic interactions between species. To do that I randomly sampled growth rates 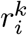 from a random uniform distribution within the range of *U*[—0.05, —0.1] for both plants and the pollinators. This particular distribution was specifically used to ensure that growth rates are negative such that mutualistic interactions are obligate. This indicates that species in a community need mutualistic interaction to maintain positive abundance. In addition, I ensured that intraspecific competition within each guild of species was strictly stronger than interspecific competition. For that *α_ii_* = 1 was fixed, and I sampled *α_ij_,i≠j* from random uniform distribution ranging from *U*[0.01, 0.05]. When ensuring intraspecific competition to be stronger than interspecific competition, mutualistic communities became feasible, provided strength in mutualistic interactions *γ*_0_ was sufficiently high (Barabás et al. 2017; Baruah 2022). This specific distribution indicates that species competition was not too strong to dictate dynamics in a mutualistic community. In addition, usage of different distribution does not impact population dynamics (Baruah 2022). These distributions were particularly used such that local dynamics became feasible and stable (Baruah 2022). In addition, the distribution of intra- and inter competition interaction strengths were within the distribution empirically measured in plant-pollinator communities (Johnson et al. 2022) or in plant communities only (Wiegand et al. 2021). In this context intraspecific competition was always found to be stronger than interspecific competition which is what I model in this study. The goal was to assess how network properties interplay with dispersal and spatial scale of interactions to impact the timing of occurrence of tipping points in mutualistic meta-networks.

Finally, dispersal among patches was constrained by the spatial scale and as well as distance among patches. Species dispersal among patches decreased exponentially as the distance among patches increased. Specifically, dispersal of species *i* from patch *k* can be written as (Thompson and Gonzalez 2017),

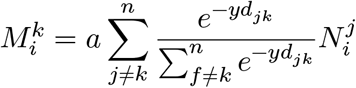

Following Thompson and Gonzalez (2017), more than one dispersal route can be taken during a particular time step, *d_jk_* is the distance between patch *j* and *k* and *y* controls the rate at which this distance impacts species dispersal, which was fixed at 0.5. Fixing it at 0.5 gives global dispersal, which meant that as metacommunity sizes became larger species dispersal did not necessarily get constrained locally, and species could in principle disperse long distance. However, the rate at which they disperse would still depend on distance between patches. Finally, *a* gives the average rate at which species disperse. Here, when we vary rate of dispersal we ensure that species dispersal remains same across guilds of species although in nature this might be species-specific. Here, three rates of species dispersal were used, *a*: 0,0.05,0.15.

### Collapse of spatial mutualistic networks

Nestedness was measured as NODF which is an acronym for nestedness metric based on overlap and decreasing fill. Nestedness of mutualistic networks that was used in this study ranged from as low as 0 to as high as 0.9, while connectance ranged from 0.08 to 0.64. Network size also ranged from as low as 8 to as high as 68.

By gradually decreasing the average mutualistic strength,*γ*_0_, globally, mutualistic networks were forced to collapse. As mutualism among guilds of species was obligate, decreasing average mutualistic strength, *γ*_0_, among species would lead to collapses of species. At a specific mutualistic strength (commonly known as threshold strength or tipping point) collapse of the entire mutualistic network would occur. Collapse of mutualistic networks thus was done by gradually decreasing *γ*_0_ from 5 to 0 in steps of 0.25. Globally, across habitat patches this scenario could be linked to changes in the climate such that phenological interaction among plants and pollinators would decrease gradually. For each value of *γ*_0_, I simulated the dynamics of the whole metacommunity for 2000 time steps. Usually, fluctuations in species density stabilize at around 1000 time points. I discarded the initial transient dynamics i.e., from *t* = 0 to *t* = 1000 and estimated equilibrium total plant and animal abundance from the last 1000 time points. Thus equilibrium network biomass was quantified as the sum of equilbrium plant and animal abundance. The extinction threshold of species in such mutualistic networks were fixed at 10^−4^. As the strength of mutualistic interactions decreased, loss of species occurred until the entire meta-mutualistic network collapsed.

Next, I determined the point of transition or the mututalistic strength at which a mutualistic network in a habitat patch *k* transitioned to a collapse state. This was quantified as when the average equilibrium meta-network biomass fell below 80 percent of its equilibrium metacommunity network biomass at maximum *γ*_0_. Once the mutualistic strength at which a network collapsed in a habitat patch *k* was determined, I evaluated the relationship of the point of transition with network topology such as connectence and network size, and how such a relationship was influenced by spatial scale of interactions.

### Predictors of temporal and spatial mutualistic network collapses

I estimated a host of spatial and temporal indicators of collapses at the community level and at the species level. For each mutualistic strength, I estimated spatial variability, regional metacommunity variability, local temporal patch variability. I quantified metacommunity biomass as the sum of all species abundance across all patches in the metacommunity. I also estimated metacommunity variability defined as the variability in abundance at the metacommunity level (see Wang and Loreau (2014) and Wang et al. (2019) for details) - 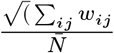, where *w_ij_* is the covariance matrix of community biomass *N_i_*(*t*) at patch *i* and 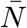 is the temporal mean of the total metacommunity biomass. Temporal variation or alpha variability was estimated as 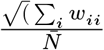 and spatial beta variability was quantified as as the variability at spatial level 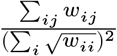 (Wang and Loreau 2014 ) for each levels of average dispersal rate, metacommunity size, and for each mutualistic threshold strength.

At the species level, I estimated standard deviation as 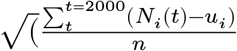 (where *n* is the number of time points and *u_i_* is the mean species abundance), and temporal autocorrelation coefficient of equilibrium species abundance for each level of mutualistic strength, for three different rates of species dispersal and metacommunity sizes. Temporal autocorrelation coefficient at first-lag and standard deviation are the classic phenomenological early warning signals that are suggested to be useful in forecasting critical transitions (Scheffer 2009; Dakos et al. 2012a) . Temporal autocorrelation coefficient at first-lag is measured as, *y*(*N*_*t*+1_) = *αy*(*N_t_*) + *σϵ_n_*, where *α* is the autocorrelation coefficient of the first-order autoregressive (AR(1)) process. *α* is close to 1 for a red-noise and close to 0 for white noise process. *α* close to 1 would indicate that the temporal abundance dynamics of a species is highly correlated and 0 would mean uncorrelated. High correlation would indicate that species is closer to a tipping point. Furthermore, I also evaluated how species degree (total unique interactions of a species) related to occurence of tipping points and the performance of species-level indicators for different levels of species dispersal and metacommunity sizes.

I wanted to evaluate whether statistical metrics measured at the species level or at the level of the community, as strength in mutualistic interaction *γ*_0_ decrease, could perform in indicating an impending critical transition. So for each level of changes in the mutualistic strength of interaction *γ*_0_ , temporal autocorrelation and standard deviation was measured at the species level, metacommunity variability was measured at the level of the metacommunity, patch variability was measured at the level of a patch, and spatial variability across habitat patches were estimated. To compare how these metrics performed in predicting collapse in relation to local network properties, I quantified Kendall’s tau correlation coefficient (Fig. 1) (Dakos et al. 2012a). Kendall’s tau rank correlation coefficient has values that range from —1 to 1 regardless of the value of the statistical metric, where 1 indicate perfect positive correlation and —1 indicate perfect negative correlation. Kendall’s tau method has been regularly used in the early warning signals literature (Scheffer 2009; Dakos et al. 2012a; Baruah et al. 2020). It indicates how strong statistical metrics such as standard deviation or autocorrelation in network biomass could increase before a network collapse. In all our simulations as we change average mutualistic strength, *γ*_0_, all networks eventually collapses. However, the question was which type of indicators would do well in predicting collapses, and what network structures or spatial scale could affect such predictability. Therefore, in our analyses, the aim was to evaluate which indicators will show strong increases in their value before eventual network collapses, and not whether one can predict accurately such collapses. To quantify the increases of such statistical indicators before network collapses, I use Kendall’s tau correlation coefficient. High positive Kendall’s tau value would indicate strong signals of network collapse. Higher the Kendall’s tau correlation coefficient, stronger is the early warning signal of network collapse and thus is of more predictive value. On the other hand, negative values would indicate false negatives, that is, a network collapsed but the above described statistical metrics failed to predict it. Next, I then evaluated how these metrics performed in relation to network properties, rate of species dispersal, and size of the metacommunity.

**Figure 1:**
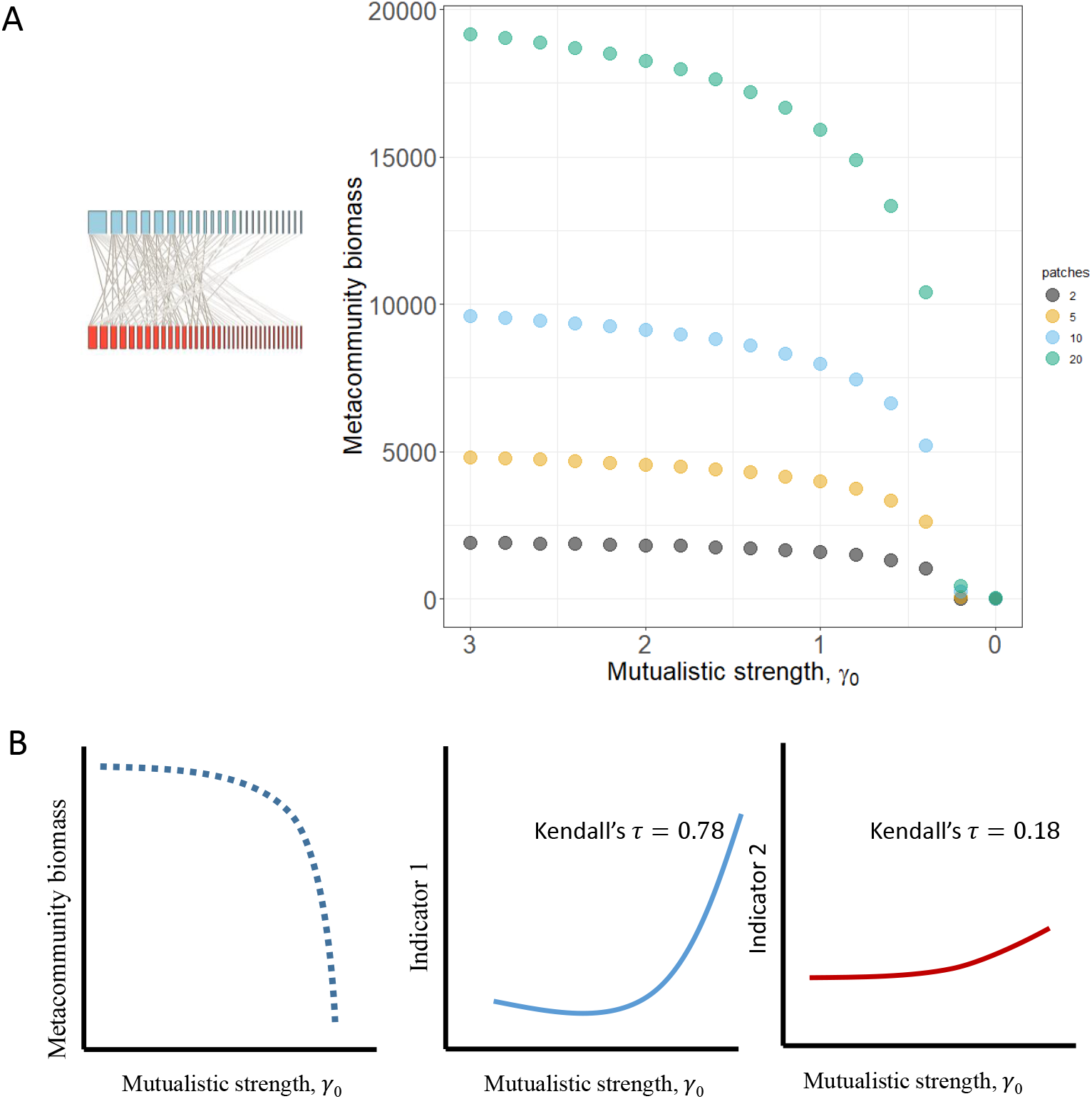
(A)Example equilibrium metacommunity abundance of a mutualistic network as mutualistic strength is decreased gradually for different sizes of metacommunity (no of patches) and rate of species dispersal (shown here for a rate of diserpsal of 0.1). (B) Shown as an example cartoon figure, as mutualistic strength decreased metacommunity biomass decreased. In addition, as biomass decreased, indicator 1 and 2 increased. This increase was quantified by Kendall tau correlation coefficient.Higher positive values would indicate stronger increase in a statistical indicator. Here indicator 1 increased strongly and thus has higher rank correlation coefficient than indicator 2, and thus is better than indicator 2.

## Results

### Point of transition and network properties

Results indicated that the mutualistic strength at which networks collapsed were determined by network size, species dispersal rate, and the size of the metacommunity. Particularly, the relationship between network size and the strength at which networks collapsed becomes strongly negative at the highest dispersal rate modulated by the size of the metacommunity (Fig. 2A). When dispersal rate was zero, the size of metacommunity on the mutualistic strength at which network collapsed remained unaffected. However, at the highest dispersal rate, smaller networks collapsed much earlier (i.e., at higher mutualistic threshold strength) than larger networks which became more evident at smaller metacommunity sizes. Connectance or nestedness didn’t have any significant impacts on the threshold at which mutualistic networks collapsed (Fig. S1). This indicated that higher species dispersal might be detrimental as networks could collapse at a much higher mutualistic threshold strength, particularly when metacommunity sizes were smaller.

**Figure 2:**
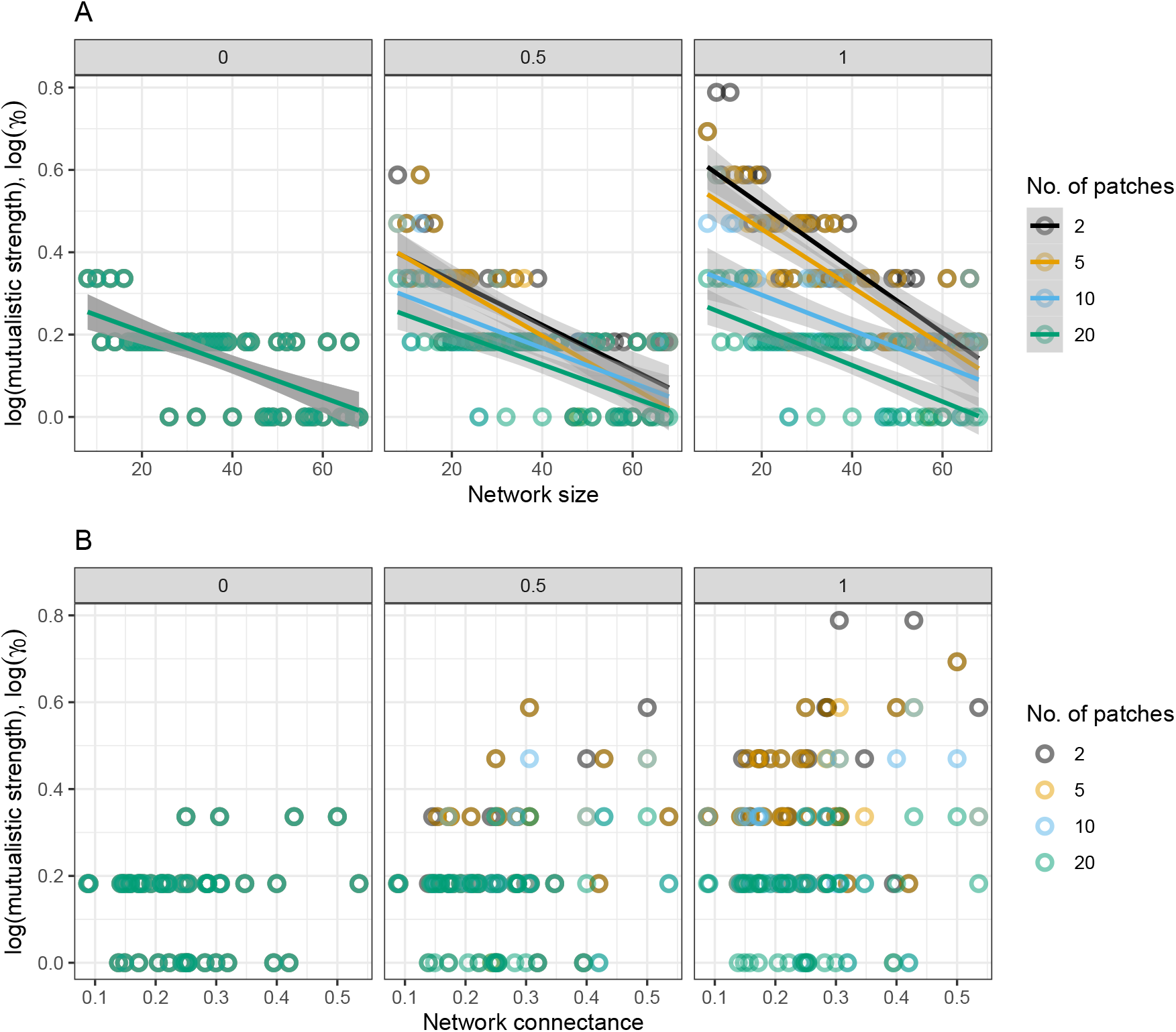
Relationship between the threshold mutualistic strength at which networks collapsed and structural properties of the network namely network size and connectance for different metacommunity sizes (2, 5, 10, 20) and three levels of species dispersal (0, 0.05, 0.15). A) Network size had a significant impact on the strength at which networks collapsed modulated mainly by the number of habitat patches and rate of species dispersal. Networks at larger metacommunities collapsed on average at a lower threshold strength that networks at smaller metacommunities. This indicated that larger metacommunities were less vulnerable to collapses at a much higher mutualistic threshold strength than smaller metacommunities. B) Network connectance did not have a significant impact on the mutualistic strength at which networks collapsed across a range of dispersal rates and metacommunity sizes.

### Indicators of transition and network properties

In an example figure of a mutualistic network (see fig. S5), I show that as metacommunity biomass collapses, indicators such as patch variability, spatial variability, metacommunity variability increases whereas indicators at the species level did not exhibit strong increases as strength in mutualistic interaction decreased. To quantify the strength of such an increased, I measure (as detailed in the methods section) the Kendall’s *τ* correlation coefficient. Such a measure of would indicate which statistical metric had the strongest increase as the metacommunity collapsed.

In case of spatial variability, when dispersal rate was zero, Kendall’s *τ* value for smaller networks were lower than larger networks as they collapsed. Since high and positive Kendall’s *τ* value captured how strong a statistical metric increased as a ecological meta-network collapsed, this particular result indicated that smaller networks exhibited less stronger increases in spatial variability as they collapsed than larger networks (Fig 3A). This result was consistent across metacommunity sizes. As dispersal increased, which networks exhibited stronger spatial variability before collapse was slightly dependent on the size of the metacommunity. Generally, larger mutualistic networks embedded in larger metacommunities exhbited strong increases in spatial variability (captured by positive values of Kendall’s *τ* ). Smaller mutualistic networks embedded in larger or smaller metacommunities exhibited less strong increases in spatial variability before collapse (Fig. 3A).

**Figure 3:**
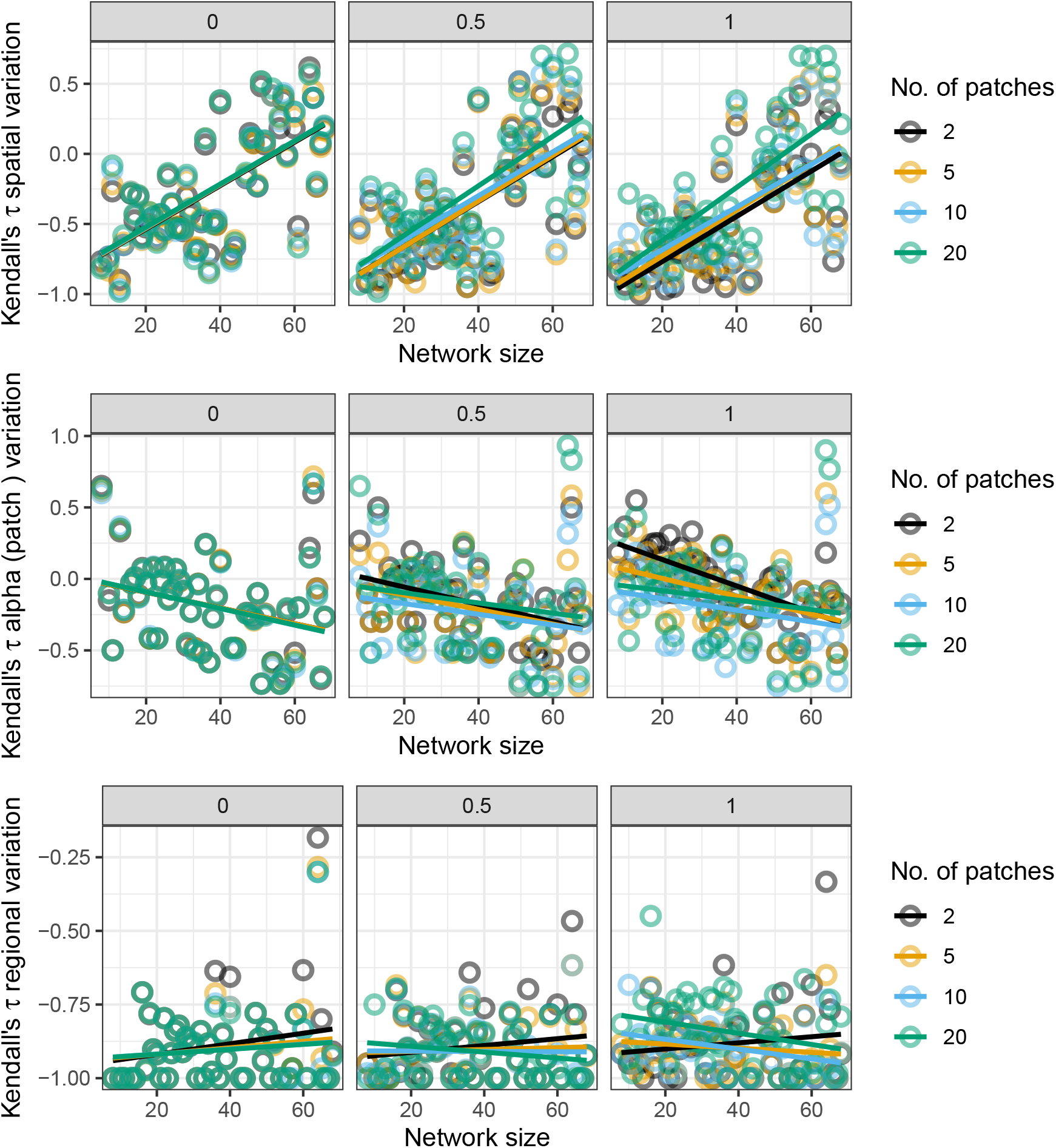
Relationship between network size and Kendalls tau coefficient of statistical metrics such as A) spatial variability, B) alpha variability, and C) metacommunity variability for three levels of species dispersal (0,0.05,0.15) and four metacommunity sizes (2, 5, 10, 20). Note that the y-axis in all these subplots indicate the strength of increases in these indicators (see figure 1C for description) as a mutualistic metacommunity collapsed. Larger positive Kendall tau values would indicated stronger increases as a mutualistic meta-network collapsed and negative values would indicate false negatives.

Patch variability also known as alpha variability was impacted negatively by the size of mutualistic networks and less impacted by species dispersal and number of habitat patches i.e. metacommunity size (Fig. 3B). Particularly, larger networks had lower Kendall’s *τ* value in comparison to smaller networks. This negative relationship however was not impacted by rate of species dispersal or the size of metacommunities significantly (Fig. 3B)

Results also indicated that when species dispersal was zero, size of mutualistic network had no impact on Kendall’s *τ* of metacommunity regional variability even at higher levels of species dispersal or for different sizes of metacommunities (Fig 3C). Since, network connectance was negatively correlated with network size, the above result remains similar except that the relationship between Kendall’s tau of metacommunity variability and connectance became slightly positive, although all values were below 0 which further suggests that the performance of metacommunity variability in forecasting meta-network collapses was rather poor (Fig. S2).

The relationship of strength of mutualistic interaction at which species collapsed as the meta-network collapsed and species degree was negative and exponential (Fig. 4A). Species which had higher number of interactions i.e., higher degree, collapsed later (at a much lower strength of mutualistic strength, *γ*_0_). This was similar across rate of species dispersal. However, size of the metacommunity had a slight impact at higher dispersal rates (Fig. 4A).

**Figure 4:**
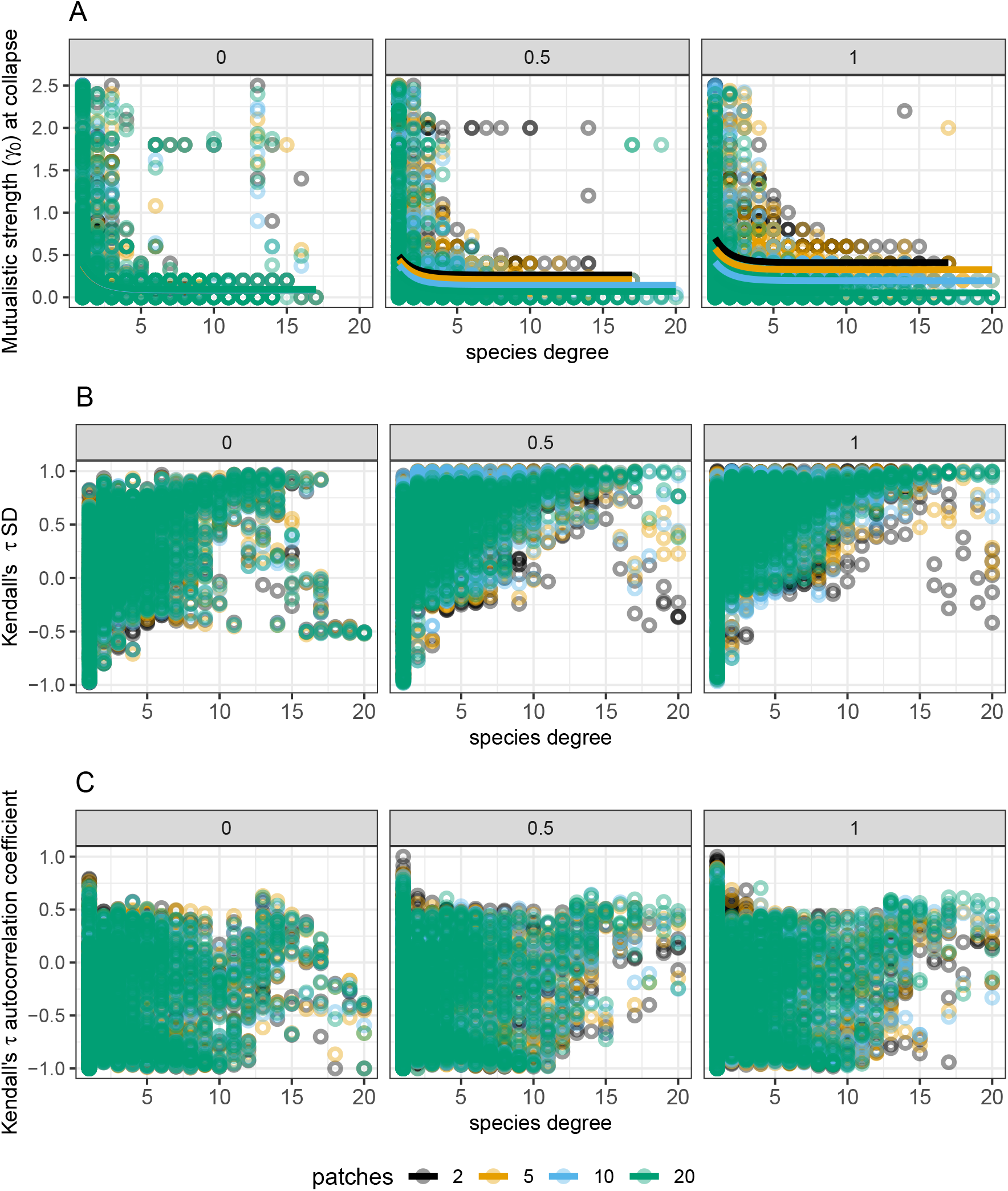
A) Strength of mutualistic interaction at which species in a network collapsed in comparison to its degree. B) Relationship between species degree and Kendall tau of SD (standard deviation) measured at the species-level and C) autocorrelation at first-lag for three levels of dispersal (0,0.5,1) and four metacommunity sizes (2, 5, 10, 20). Note that the y-axis in all these subplots indicate the strength of increases in leading indicators as a mutualistic metacommunity collapsed. Higher and positive Kendall tau would indicated stronger increases as a mutualistic meta-network

Strength in warning signals such as standard deviation measured at the species-level was related to species degree, and less slightly on rate of species dispersal and the size of the metacommunity. Strength of SD, measured as the Kendall’s *τ* correlation coefficient, increased as species degree increased indicating that species which have a larger number of unique interactions will display stronger increases in SD as the network collapses due to changes in the environment that weakens mutualistic interactions. Strength in autocorrelation, however, did not relate to species degree or species dispersal. In addition, autocorrelation at first-lag, or standard deviation measured at the species level was slightly impacted by topological properties such as nestedness or network size but was not impacted by species dispersal or size of the metacommunity (Fig. 4A, S4).

## Discussion

Climate change can cause shifts in interaction strengths that could lead to mismatches in species interactions thereby causing negative impacts on species biomass and diversity (Revilla et al. 2015; Thompson and Gonzalez 2017). In mutualistic networks, positive interactions between groups of species leads to a positive-feedback mechanism that promotes the occurrence of sudden collapses once environmental change causes interaction strengths to fall below a certain threshold (Lever et al. 2014; Kéfi et al. 2016; Baruah 2022). Although, studies have tried to understand how such networks are robust to habitat destruction, and changes in phenology, little is known about how the architecture of such mutualistic networks at the spatial scale could influence the timing of such collapses. Here, I show that species disperal rate, size of the metacommunity, and network architecture has significant impacts on the threshold at which a mutualistic meta-network collapses. Furthermore, performance of statistical metrics that could be used to forecast such unwanted collapses, such as metacommunitiy variability, alpha variability or spatial variability, was not only dependent on the architecture of the mutualistic meta-network but also on species dispersal rate, and the size of the metacommunity.

When species dispersal rate was zero, all mutualistic networks distributed across a spatial scale could be coalesed to independent local communities. Effectively, metacommunity size i.e., number of habitat patches in the metacommunity, thus, had no impact when a network collapsed. This was obvious as species dipsersal was zero and spatial insurance provided by species dispersing across the landscape was effectively nil (Loreau et al. 2003). As a result, when strength of mutualism decreased, networks collapsed and metacommunity size did not have any impact on the threshold strength at which these networks collapsed. There was, however, a slight negative relationship between size of the network and the mutualistic threshold at which such networks collapsed (Fig. 2A). Particularly, larger networks collapsed later than smaller networks. Earlier studies have indicated that feasibility of mutualistic networks increases as network size or nestedness increases (Lever et al. 2014; Baruah 2022). This was intuitive as larger networks have larger number of species which effectively increases network biomass and persistence of the network as a whole.

As species dispersal increases, the effect of metacommunity size became more evident. At high species dispersal, the relationship between network size and the mutualistic threshold strength at which networks collapsed became more negative. Smaller networks collapsed much earlier (higher average mutualistic strength, ($ _0 $) at higher dispersal rate than larger networks. This particular result was evident at smaller metacommunity sizes (Fig. 2A). At smaller metacommunities, for instance a metacommunity with two habitat patches, species were able to easily disperse to the only other habitat and thereby homogenise communities. Higher dispersal, as indicated in previous studies (Loreau et al. 2003) , tend to lead to lower species richness. As such, at high rate of species dispersal and in smaller metacommunities, smaller networks lost species faster at a lower mutualistic threshold than larger networks. In such mutualistic metacommunities species were modelled to be obligate mutualists, which indicated that species were dependent solely on mutualistic interactions for maintaining positive growth rate. As species dispersal increased, a stronger average mutualistic interaction was required for to maintain an overall positive growth rate locally. As a consequence, at very high dispersal rates patches became homogenized, particularly those that had a fewer number of patches, and hence, a much stronger mutualistic strength on average was required to maintain positive abundance. This particular phenomena did not become an issue for larger metacommunities, for instance a metacommunity with more than five habitat patches and for larger networks (Loreau et al. 2003). Larger networks, in addition, harbored higher number of species and as such network collapse occurred at a much lower mutualistic threshold strength than smaller networks. Results, therefore, indicate that in order to preserve biodiversity, it is thus imperative to take into account both the sizes of communities (topological features) as well as the spatial scale of species interactions (Thompson and Gonzalez 2017).

Much work had been done in identifying statistical signals that could forecast unwanted critical transitions. These statistical signals are phenomenological in nature and could be easily identified from state-based time series data such as abundance or biomass (Dakos et al. 2012b, a; Scheffer et al. 2012; Clements and Ozgul 2016; Baruah et al. 2019). However, such signals and their efficacy have been questioned recently both in numerical simulations studies (Hastings and Wysham 2010; Clements et al. 2015a; Baruah et al. 2020) and experimental studies (Wilkinson et al. 2018; Baruah et al. 2021). Their has been a few application on multi-species communities (Dakos 2017; Patterson et al. 2021; Baruah et al. 2022). Here, I evaluated how the efficacy of commonly used statistical signals measured at the species level (autocorrelation at first-lag, standard deviation), community level (patch variability, spatial variability) and at the metacommunity level (regional variability) perform in forecasting collapses in relation to network topology and spatial scale of interactions. Interestingly, species level metrics had a positive relationship with network architecture such as nestedness, but was not impacted by either dispersal rate of species or the size of the metacommunity (Fig. S4). Community and metacommunity-based signals such as alpha variability, spatial variability also increased as mutualistic strength decreased. Smaller networks in metacommunities collapsed earlier and such networks also exhibited weaker increases in spatial variability across different species dispersal rates (Fig. 3A-C). This indicated that spatial variability performed poorly in forecasting the collapse of smaller networks. Alpha variability, however, exhibited an opposite but less steep increase of a pattern. Thus, when global changes weaken mutualistic interactions thereby leading to loss of biodiversity, indicators measured at the community and metacommunity level could potentially inform instability. However, smaller networks could exhibit less strong signals in contrast to larger networks across a range of metacommunity sizes. This analysis indicated that these indicators and their performance when estimated at the community or metacommunity level would be slightly dependent on species dispersal, network architecture, and how large the metacommunity was.

Changes in the environment can weaken species interactions to the point of biodiversity collapse. Recovering lost ecosystem functions and processes is not easy as demographic information of species are also lost as biodiversity collapses [(Scheffer 2009; Link and Watson 2019). In addition, most ecological systems can exhibit a phenomenon called hysteresis whereby even if the original stable environmental conditions are reverted, the ecological system might still not recover. There are tools that have been thus develop to forecast such unwanted transitions (Wissel 1984; Dakos et al. 2012a). When an ecological network collapses both at the local as well as at the spatial scale, signals of global meta-network instability could also be manifested in species in an ecological community. It is relatively unknown, however, which species could exhibit signals of instability, but see (Dakos 2017; Patterson et al. 2021; Baruah et al. 2022). Here, the strength at which species collapsed was dependent on species degree in a non-linear way. The relationship between species degree and mutualistic threshold strength at which species of a network collapsed was negative exponential. This result meant that species having a higher degree of interactions were more resistant to changes in mutualistic interaction strength than species with lower number of interactions. However, the difference in mutualistic strength at which, say, species with degree three collapsed and a species with degree eight collapsed was negligible (Fig. 4A). Only species that have the fewest number of interactions (less than three) collapsed at a much lower mutualistic strength. In addition, the species that exhibited strong increases in standard deviation as networks collapsed, were the ones which had on average a larger number of interactions (Fig. 4B). The relationship between species degree and strength of species SD was non-linear, indicating that species with moderate to high number of interactions would show the strongest increase as networks collapsed. Species that had a relatively larger number of mutualistic interactions, benefited from higher positive growth rate which resulted in stronger increases in standard deviation as networks collapsed than those species that had fewer interactions. However, as shown in Dakos and Bascompte (2014), and also here, species that have fewer interactions would collapse earlier and might not exhibit as strong an increase in standard deviation as networks collapsed. This indicates that, although, one could potentially label species that have higher number of interactions as “indicator species”, that, however, would not help in informing instability of “specialist species” as they generally collapsed much earlier (Fig. 4A). Nevertheless, these results indicate that standard deviation measured at the species level could potentially increase as mutualistic interactions weaken and more so for species with a moderate to high number of interactions. Autocorrelation at first-lag also increased as interactions weakened, but its relation with species degree was unclear.

Our ability to detect an impending transition is dependent on monitoring dynamics at the vicinity of the transition. However, monitoring population dynamics required intense temporal as well as spatial sampling and could potentially impact forecasting of abrupt transitions (Arkilanian et al. 2020; Bruel and White 2021) . Many other factors could also impact prediction of such transitions including but not limited to long transients and stochasticity (Hastings and Wysham 2010; Hastings et al. 2018). There has been quite a debate on the appropriate set of methods or tools that could be used to predict future biodiversity states. Studies have indicated that in addition to monitoring dynamics of species abundance, phenotypic traits should be monitored as well. Including information from phenotypic dynamics such as body size could improve forecasts of biodiversity collapse (Clements and Ozgul 2016), although such an accurate forecast of biodiversity collapse is dependent on the type of environmental perturbation and the type of interactions prevalent in the community (Baruah et al. 2022). When changes in the environment impact communities both locally and spatially alike, ecological networks could abruptly collapse. However, when such networks collapses depends on the average rate of species dispersal, with higher species dispersal and smaller metacommunities causing meta-networks to collapse at a much higher mutualistic strength. In addition, collapses of mutualistic meta-networks was also dependent largely on the topology of the network, with smaller networks collapsing at a higher mutualistic strength than larger networks. These results are pertinent for conservation efforts particularly because it points to the fact that ecological networks at smaller metacommunities are as vulnerable or if not more than meta-networks at a much larger spatial scale. When assessing whether communities are vulnerable to changes in the environment, the results from this research indicates that it is very pertinent to look at factors beyond species demographic rates, that includes the scale of spatial interactions, and topology of local ecological communities.

## Author contributions

GB conceptualized the study, did the analysis and wrote the manuscript.

## Data availability

All the data and scripts will be archived in Zenodo upon acceptance. The 101empirical networks were accessed from www.web-of-life.es.

The paper was written in rmarkdown format and files, data, and R scripts to reproduce the paper is also located at the github repository : https://github.com/GauravKBaruah/spatial_tipping_points

## Supplementary document

**Figure S1:**
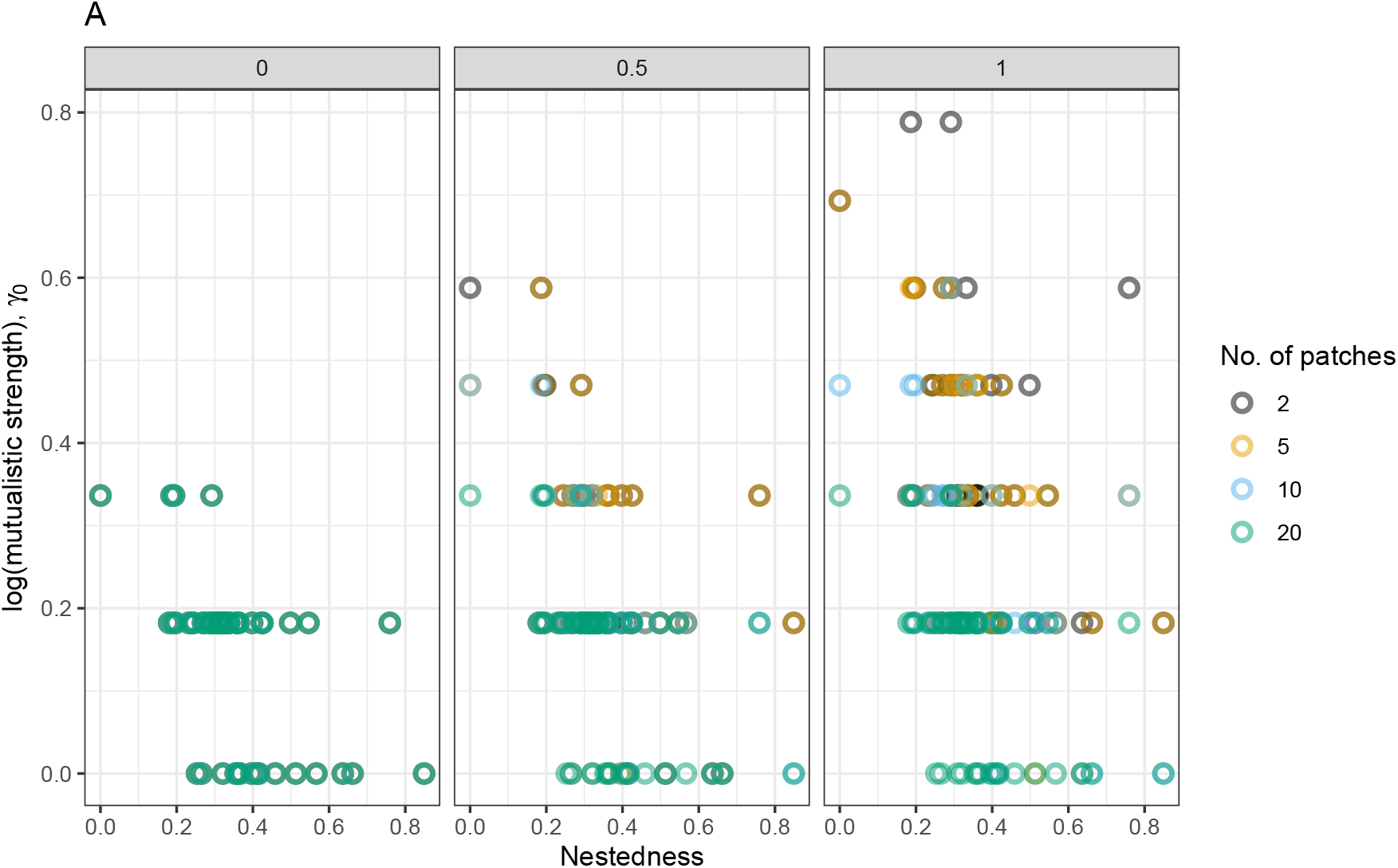
Relationship between the threshold mutualistic strength at which networks collapsed and structural properties of the network namely nestedness (NODF).

**Figure S2:**
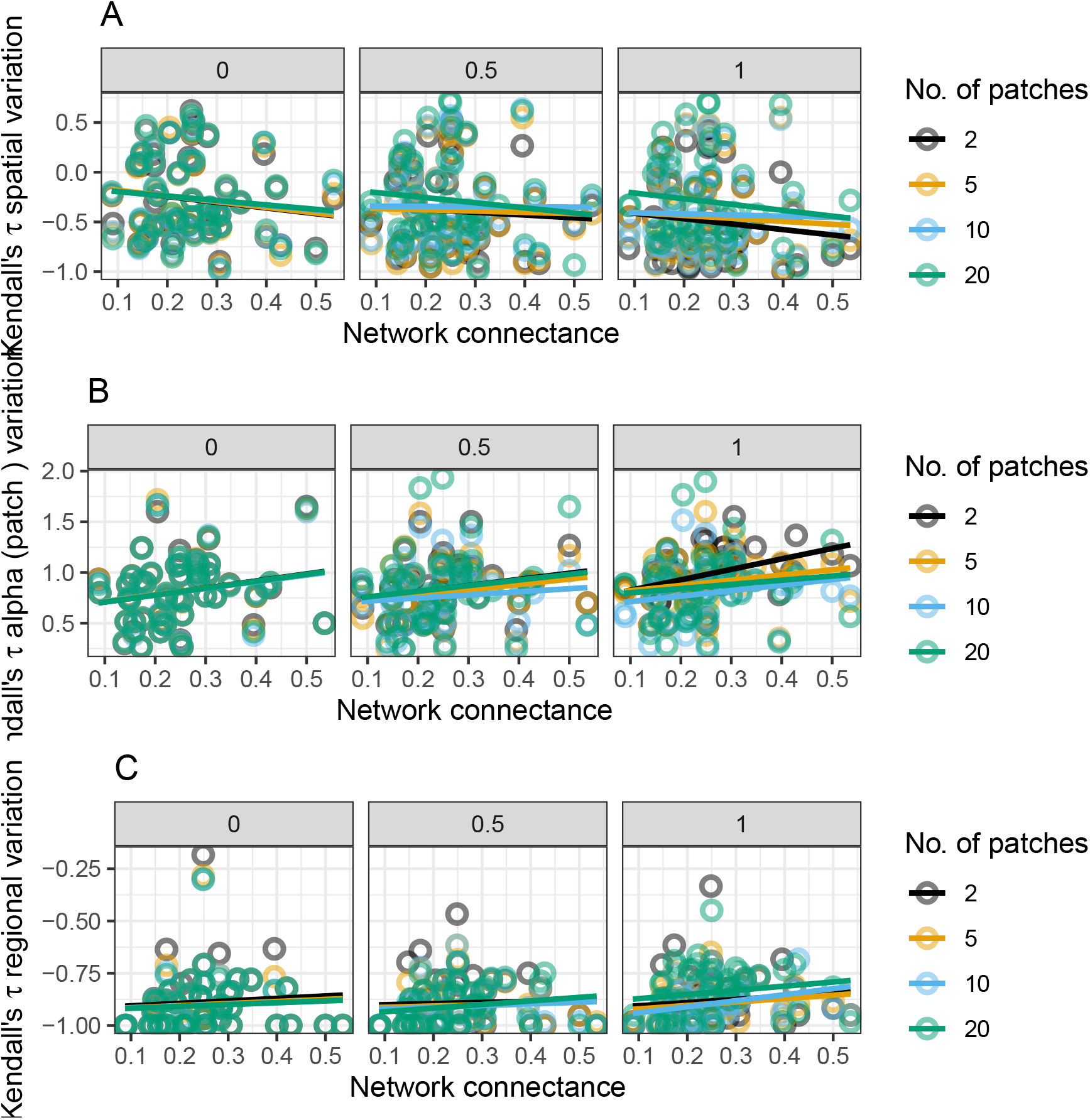
Relationship between network connectance and Kendall tau coefficient of statistical metrics such as A) spatial variability, B) Alpha variability, and C) metacommunity variability for three levels of species dispersal (0,0.05,0.15) and four metacommunity sizes (2, 5, 10, 20). Note that the y-axis in all these subplots indicate the strength of increases in these indicators (measured as Kendall tau coefficient, see figure 1C for description) as a mutualistic metacommunity collapsed. Higher positive values of Kendall tau coefficient would indicate stronger increases as a mutualistic meta-network collapsed.

**Figure S3:**
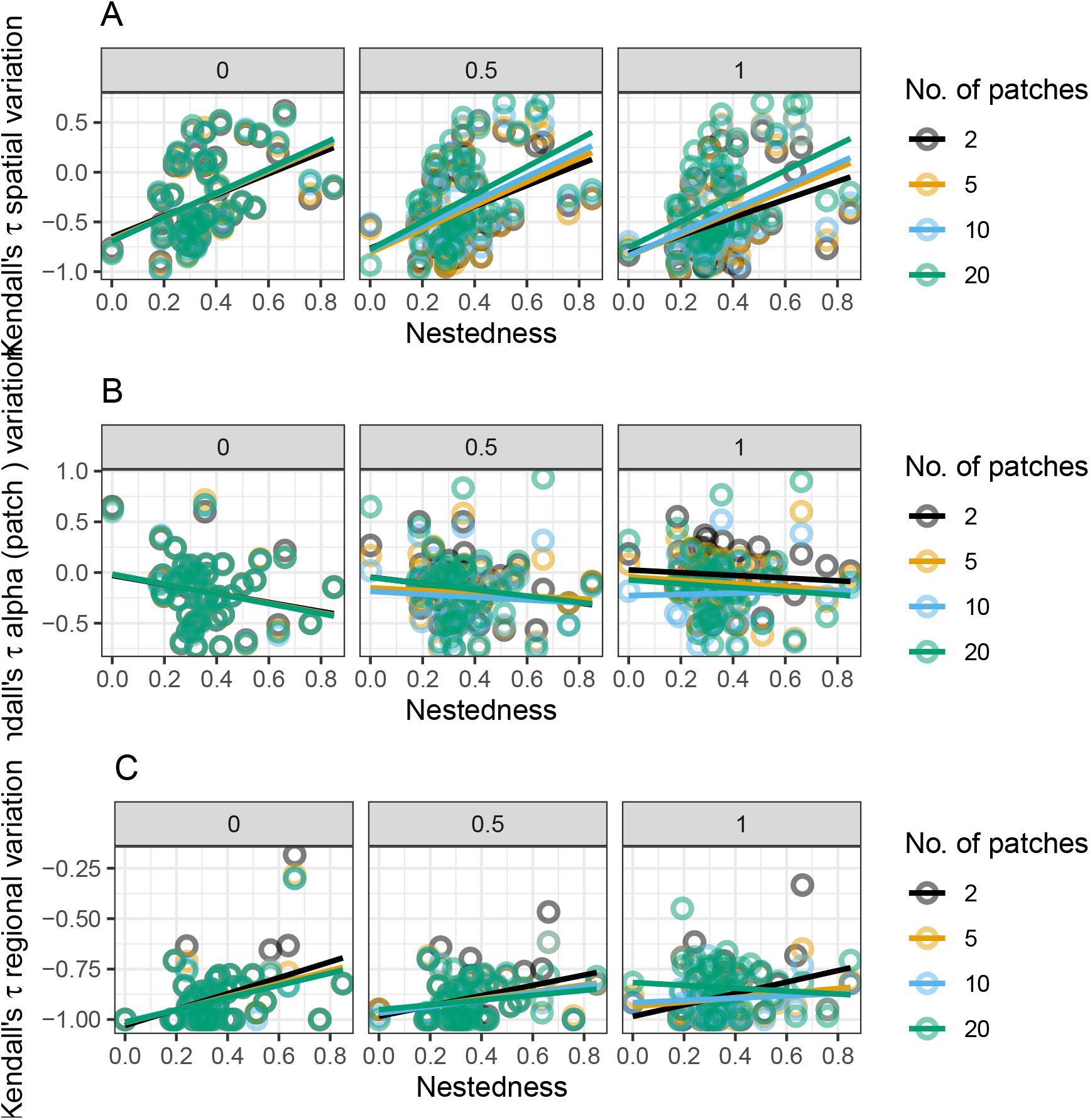
Relationship between network nestedness (NODF) and Kendall tau coefficient of statistical metrics such as A) spatial variability, B) Alpha variability, and C) metacommunity variability for three levels of species dispersal (0,0.05,0.15) and four metacommunity sizes (2, 5, 10, 20). Note that the y-axis in all these subplots indicate the strength of increases in these indicators (measured as Kendall tau coefficient, see figure 1C for description) as a mutualistic metacommunity collapsed. Higher positive values of Kendall tau coefficient would indicate stronger increases as a mutualistic meta-network collapsed.

**Figure S4:**
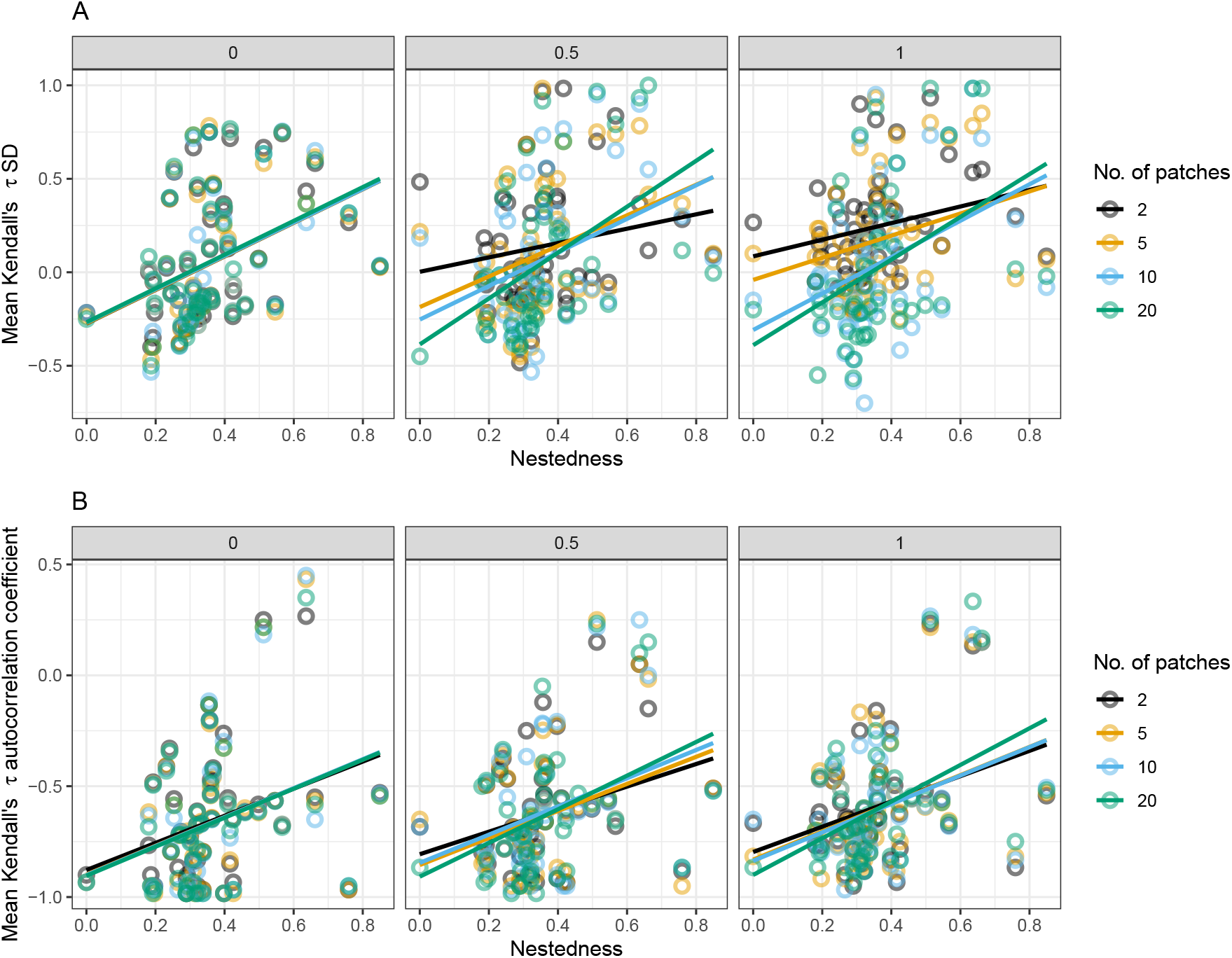
Relationship between network nestedness (NODF) and Kendall tau coefficient of statistical metrics such as A) species SD, B) Autocorrelation coefficient at first-lag, for three levels of species dispersal (0,0.05,0.15) and four metacommunity sizes (2, 5, 10, 20). Note that the y-axis in all these subplots indicate the strength of increases in these indicators (measured as Kendall tau coefficient, see figure 1C for description) as a mutualistic metacommunity collapsed. Higher positive values of Kendall tau coefficient would indicate stronger increases as a mutualistic metanetwork collapsed.

**Figure S5:**
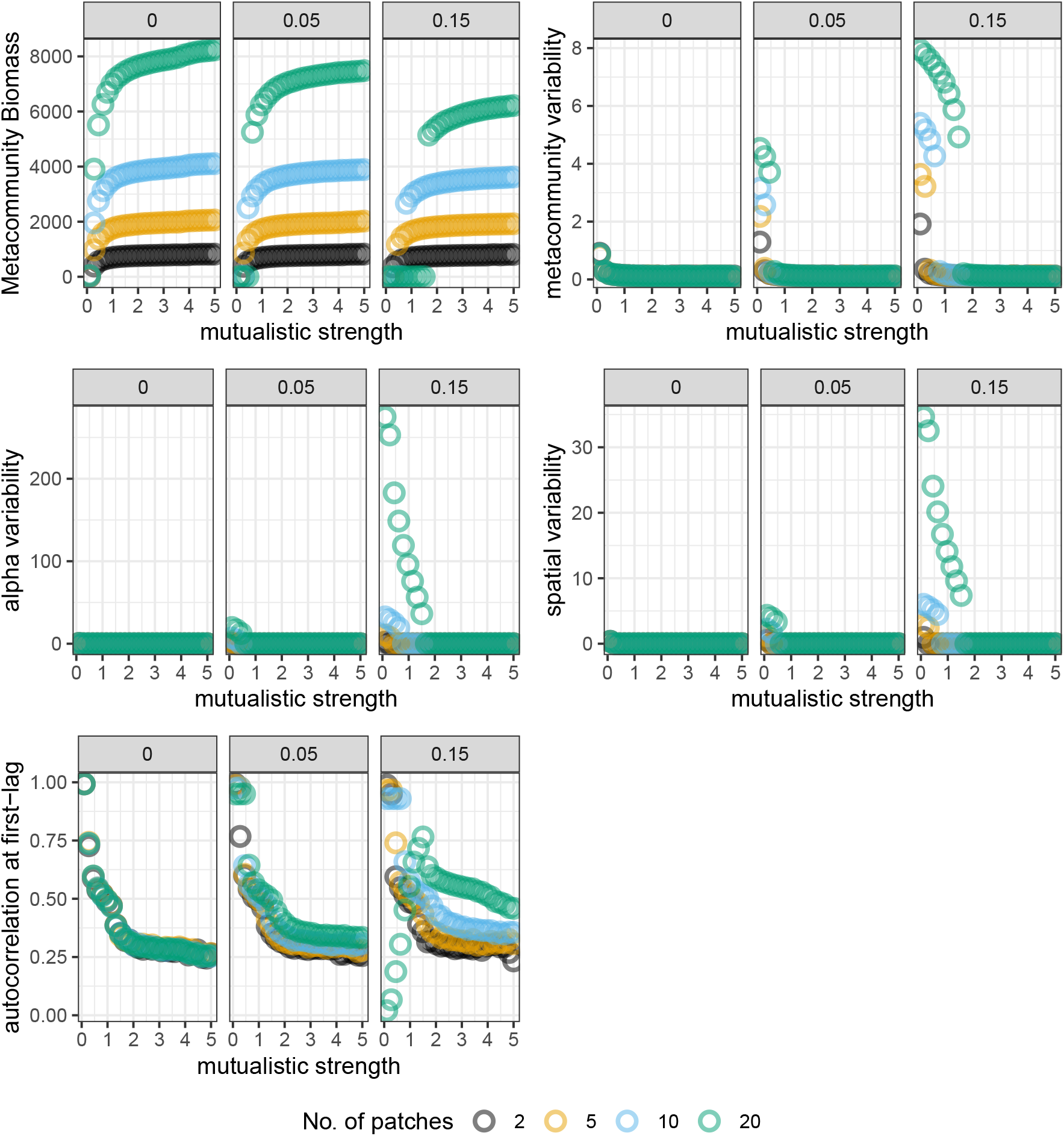
Meta-network collapse and statistical metrics that increased as mutualistic strenght decreased for three rates of dispersal, four metacommunity size. A) Example meta-network collapse as mutualistic strength decreased. (B) metacommunity variability as mutualistic strength decreased. (C) alpha variability as mutualistic strength decreased. (D) Spatial variability as mutualistic strength decreased, (E) species autocorrelation as mutualistic strength decreased.

**Figure S6:**
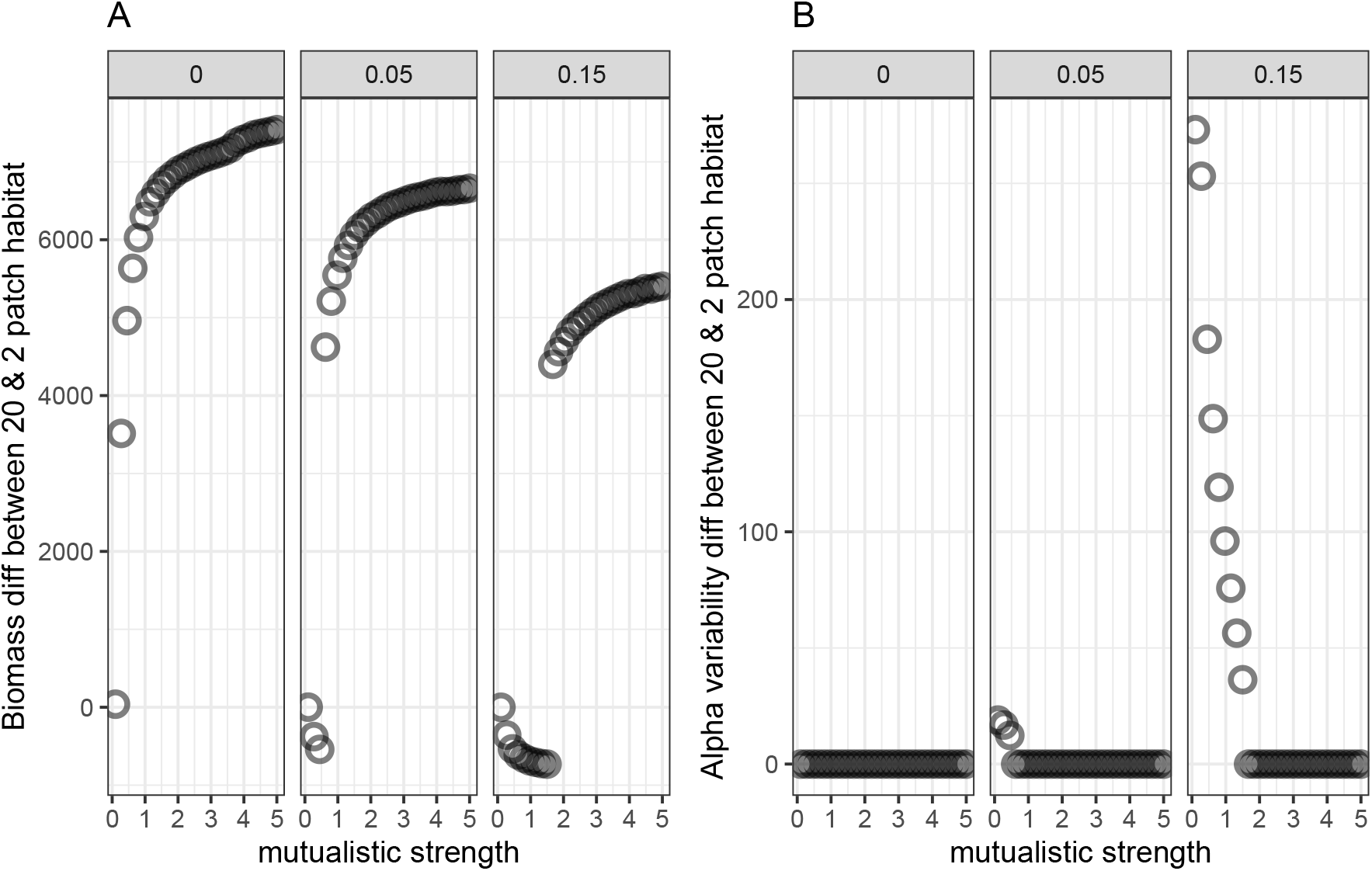
A) Metacommunity biomass difference of mutualistic network between 20 patch habitat and 2 patch habitat for three range of dispersal rates (0, 0.05, 0.15) and for a range of mutualistic strenght. B) Difference in alpha variability between 20 patch habitat and 2 patch habitat. Shown here is the result for one mutualistic meta network. To be noted here that ”diff” means difference.

## Notes

### Competing Interest Statement

The authors have declared no competing interest.

### Summary of Updates

Updated new analysis and figures

